# Sizing up spotted lanternfly nymphs for instar determination and growth allometry

**DOI:** 10.1101/2022.03.07.483361

**Authors:** Theodore Bien, Benjamin H. Alexander, Eva White, S. Tonia Hsieh, Suzanne Amador Kane

## Abstract

A major ongoing research effort seeks to understand the behavior, ecology and control of the spotted lanternfly (SLF) (*Lycorma delicatula*), a highly invasive pest in the U.S. and South Korea. These insects undergo four nymphal stages (instars) before reaching adulthood, and appear to shift host plant preferences, feeding, dispersal and survival patterns, anti-predator behaviors, and response to traps and chemical controls with each stage. However, categorizing SLF life stage is challenging for the first three instars, which have the same coloration and shape, because no comprehensive datasets exist. We present a dataset of body mass and length for SLF nymphs throughout two growing seasons and compare our results with previously-published ranges of instar body lengths. An analysis using two clustering methods revealed that 1^st^-3^rd^ instar body mass and length fell into distinct clusters that were consistent between years, supporting using these metrics to stage nymphs during a single growing season. However, the ranges for 2^nd^-4th instars were not consistent between our results and those from earlier studies for diverse locations. The scaling of SLF nymph body mass with body length varied between isometry (constant shape) and positive allometry (growing faster than predicted by isometry) in the two years studied. Using previously published data, we also found that SLF nymph adhesive footpad area varies in direct proportion to weight, suggesting that footpad adhesion is independent of nymphal stage, while their tarsal claws display positive allometry and hence disproportionately increasing grasp (mechanical adhesion). By contrast, mouthpart dimensions are weakly correlated with body length, consistent with predictions that these features should reflect preferred host plant characteristics rather than body size. We recommend future studies use the body mass vs length growth curve as a fitness benchmark to study how SLF instar development depends on factors such as hatch date, host plant, temperature, and geographic location, to further understanding of life history patterns that help prevent further spread of this invasive insect.

## Introduction

The spotted lanternfly (SLF), *Lycorma delicatula* White (Hemiptera: Fulgoridae) is a planthopper native to south Asia that has become a highly invasive pest in the U.S. and South Korea. SLFs feed intensively on phloem from a wide variety of trees and other plants, stressing the hosts as well as promoting the growth of sooty mold (1). Because SLFs threaten significant economic damage to agricultural crops, native trees, and landscape plants, a large ongoing research effort seeks to understand their development, physiology, behavior and ecology to inform methods for mitigation and control (2–4). In this study, we discuss how clustering methods can be applied to measurements of the body mass and size of immature SLFs (nymphs) in order to improve the determination of SLF life stage and to study the scaling of previously published SLF footpart and mouthpart dimensions (5) with body size. We begin by explaining how these issues are relevant to a wide variety of topics in SLF research.

After emerging, SLFs develop through five life stages separated by molting: four nymphal instars and the much larger and winged adult stage. The 4th instars are readily identified by their distinctive red, black and white spotted coloration. However, the first through third instars have similar black-and-white-spotted coloration and overall body morphology (Fig 1). Many studies of SLF behavior, ecology, and phenology have relied on determination of the nymphal stage (instar determination) in order to track how life stage influences ecology and choice of host plants (1,2), dispersal patterns (3–6), locomotor behaviors such as climbing and jumping (7,8), phenology and activity (9), spectral preferences (10), attraction to chemicals (11), and effectiveness of various trapping methods (12). Thus, instar determination methods for identifying the life stage of a given specimen collected in the field are useful and important in many contexts. Several previous studies have shown how a detailed microscopic examination can reveal foot, mouth part and antenna morphological changes during development (7,13,14), providing information of great utility for how these factors influence feeding, adhesion and locomotion throughout the insect’s life cycle. In practice, the life stages of the first three SLF instars have been estimated in many studies using overall body dimensions readily measured in the field, along with previously published size ranges for each instar.

**Fig 1.**
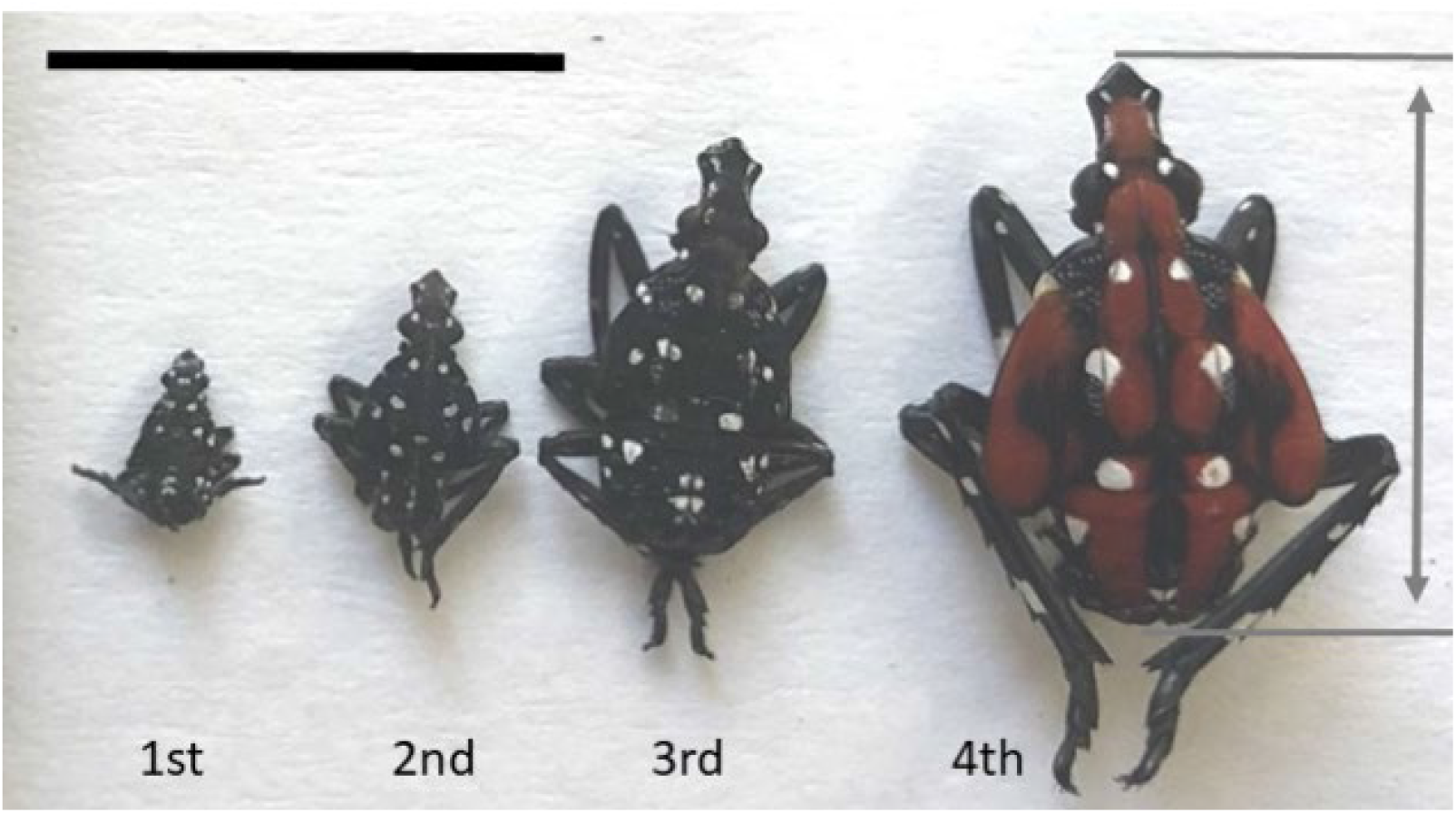
Photograph of 1st, 2nd, 3rd, and 4th instar spotted lanternfly nymphs. Double-headed arrow shows the definition of body length, L. (scale bar = 10 mm). For many insects and other arthropods, instar determination (i.e., identifying the life stage of nymphs collected in the field) relies on the observation that major changes in body dimensions occur primarily when the exoskeleton is shed upon molting (18). Ideally, this involves directly measuring the frequency distribution of one or other metrics of exoskeleton size for each instar using laboratory-reared specimens with known molting status (e.g., based on molted head capsule dimensions) (22). However, instar determination should be possible without knowledge of molting status if the number of developmental stages is known in advance, the morphometric data is uniformly sampled across all life stages, and its frequency distribution is partitioned into distinct clusters (23). The last approach is especially useful for SLFs, which have proven challenging to raise in the laboratory so that life stage can be directly monitored (20,21), and which we have observed to have flaccid cast exoskeletons that do not provide useful sizing information after molting.

In spite of this growing interest, only a few previous studies have reported measured data for the ranges of body lengths corresponding to each nymphal life stage for use in instar determination, and none have reported body mass. (Table 1, Fig 1) The earliest study reported only mean body lengths for each stage in China (15). Park and colleagues (16) measured body lengths for 1st through 4th instars in South Korea, although it was not stated whether specimens used for measurements were raised in the laboratory with known life stage or collected from the wild and the instar stage estimated from size. Jang et al. (10) reported only body lengths for just 2nd instars captured in the field in South Korea. Dara et al. (17) reported the ranges of body lengths measured for 1st through 4th instar nymphs collected in Pennsylvania. None of these previous studies provided statistical data to guide the classification of new datasets for instar determination. Furthermore, studies have shown that the size ranges for each instar can depend on factors such as date of emergence, diet, host plants, temperatures, and environment (e.g., laboratory vs field-raised) (18). Indeed, prior research has indicated that SLF nymphs develop and survive differently when reared with different diets in the field (19,20), at different temperatures (21), and artificial conditions (i.e., enclosures or laboratory conditions) (1,16), but these studies did not consider how these factors affected instar morphometrics.

**Table 1.** Body length (mm) of spotted lanternfly nymphs from this study and earlier work. N = number of specimens (not reported in (15)).

| Body length (mm) |  |  |  |  |  |  |
| --- | --- | --- | --- | --- | --- | --- |
| Life stage | This study 2022<br>mean ± SD | This study 2021<br>mean ± SD | Dara et al., 2015 (17)<br>[range] | Jang et al., 2013 (10)<br>mean ± SD | Park et al., 2009 (16)<br>mean ± SE | Zhou, 1992 (15)<br>mean |
| 1<sup>st</sup> instar | 4.32 ± 0.35<br>N = 49 | 4.24 ± 0.24<br>N = 54 | [3.6, 4.4]<br>N = 12 | Not measured | 3.9 ± 0.2<br>N = 43 | 4 |
| 2<sup>nd</sup> instar | 6.73 ± 0.38<br>N = 88 | 6.67 ± 0.48<br>N = 30 | [5.1, 6.4]<br>N = 10 | 6.0 ± 0.5<br>N = 17 | 5.7 ± 0.7<br>N = 62 | 7 |
| 3<sup>rd</sup> instar | 9.50 ± 0.54<br>N = 49 | 9.30 ± 0.72<br>N = 61 | [6.9, 9.4]<br>N = 12 | 8.6 ± 0.5<br>N not given | 8.9 ± 0.4<br>N = 23 | 10 |
| 4<sup>th</sup> instar | 12.28 ± 0.60<br>N = 40 | 11.74 ± .75<br>N = 49 | [10.9, 14.8]<br>N = 10 | 11.3 ± 0.7<br>N not given | 11.6 ± 2.3<br>N = 37 | 13 |

In this study, we report measurements of mass and body length for spotted lanternfly nymphs along with clustering results for these specimens. By comparing these results with those from four previous studies, we explore the variation in SLF nymph size distributions reported thus far. A growth (ontogenetic) allometric analysis was also performed to identify possible adaptations for feeding morphology and biomechanics. Body mass has been found to scale as a power law of body length (i.e., M = L^c^) for a wide range of insect and other arthropod taxa with scaling exponents c that vary from < 1 to 3 (24), where c = 3 corresponds to isometric growth (geometrical similarity; maintaining a constant shape), c > 3 to positive allometry (i.e., growing faster than predicted by isometry) and c < 3 to negative allometric growth (i.e., growing more slowly than predicted by isometry). We used our data to determine the ontogenetic scaling regime of SLF nymph body mass, and use previously-published morphometric data to determine how the dimensions of foot and mouth body parts scale with overall body size. We interpret these results in relation to SLF behavior and ecology, and suggest ways these methods can be applied in future work.

## Methods

### Insect collection and morphometrics

Healthy, intact SLF nymphs were collected in the field from *Ailanthus altissima* trees and wild grape vines (*Vitis spp*.) in southeastern Pennsylvania (40°00’30.2”N 75°18’22.0”W) from May through July, 2021 and 2022, corresponding to 1st instar emergence until it was difficult to find 4^th^ instars (note this was a different date in the two years). In both years, we measured all specimens collected from the field site using an insect net to avoid sampling bias. Details on the collection timeline and number of specimens collected, which corresponded to multiple samples per instar (2-4 weeks/stage in 2021; 4-5 weeks/stage in 2022) are given in S1 Appendix. A total of N = 194 (2021) and N = 226 (2022) specimens were collected across all nymphal life stages (Table 1). Because SLF are identified as an invasive species in Pennsylvania, all specimens were euthanized by freezing (25). Morphometric data were measured post-mortem after thawing for 15 min to preserve tissue hydration and morphology using an analytical balance (Explorer, Ohaus, Parsippany, NJ US) to measure mass, M (accuracy ± 0.4 mg). Body length, L, was defined as distance between the anterior end of the head to the posterior end of the abdomen; it was measured using ImageJ (26) to ± 0.05 mm from digital micrograph images of specimens lying flat on their dorsal or ventral surfaces with a scale bar in the same plane. Micrographs taken from a variety of perspectives indicated that the effect of specimen orientation was ≤ 4% of body length and hence ≤ measurement uncertainty.

### Comparison with other studies

We performed Google Scholar searches using the keywords spotted lanternfly and *Lycorma delicatula*, yielding over 600 references. Approximately 100 papers that directly studied SLFs were used to perform repeated forward and reverse citation searches to find morphometric data for spotted lanternfly nymphs. This resulted in the identification of four papers with additional values of body length (10,15–17). We also found one study that reported adult SLF body length and mass, which agreed with our own observations (27). One study reported morphometric data for footpart and mouthpart dimensions (14); here we consider their values for the tarsal claw tip-to-tip distance (their TCT), the area, A_adh_, of the arolium (adhesive pad) estimated from their arolium morphometric data, as well as the lengths of the labium, L_L_, and stylet, L_S_, which is used to pierce plant surfaces for feeding. (See S2 Appendix for more information about footpart measures.)

### Data analysis and statistics

Data analysis was performed using MATLAB version R2021a with the curve fitting and statistics and machine learning toolboxes (Mathworks, Natick MA USA); MATLAB functions are referred to using italicized names. Results are reported as mean [95% CI] unless indicated otherwise. All data and code required to reproduce all results and figures discussed here are accessible at https://doi.org/10.6084/m9.figshare.19287389.

All 4th instars were identified by their red, black and white coloring. Length and mass data for all specimens with black and white coloration consistent with 1st through 3rd instars were standardized before clustering by converting them into z-scores (i.e., zero mean and standard deviation = 1). For the first clustering method, the standardized data were fit to a three component Gaussian Mixture Model using *fitgmdist* (covariance type = full, shared covariance = false), then sorted into three components (clusters) using *cluster* in MATLAB to reflect the known number of instar stages in the dataset. We also partitioned only the length data for the first, second and third instars into three clusters using the Gaussian Mixture Model and *kmeans* for k-means clustering.

We next analyzed the mean lengths for each estimated instar to determine whether they follow Dyar’s Rule, the observation that instar body dimensions increase in size by a constant growth ratio between successive instars (28) This implies that log L_j_ = j × log G + log L_0_, where j = instar number, L _j_ = mean body length of the jth instar, and G = growth ratio = L_j+1_ / L _j_ (18). We used simple ordinary linear regression (MATLAB *fitlm*) to fit body length data from this study and previous work vs estimated instar number; we also computed as goodness-of-fit measures the F-statistic and p-value for significance testing (alpha = 0.05; null hypothesis no dependence on the independent variable), and R-squared. We used MATLAB *confint* to find 95% CI of all fit parameters. To fit body length data from previous studies, we used either means or middle of the quoted range, depending on which statistics were provided. (Table 1)

To determine the power law dependence of body mass on body length (i.e., M = a L^c^), we first log-transformed the M and L data, and then used simple ordinary linear regression to fit to log M = a + c log L, as described above. The fitted slope (the scaling exponent, c) was used to determine whether these data were consistent with the null hypothesis of isometric scaling (i.e., c = 3), or instead with positive or negative allometry. (See full results in S4 Appendix).

We performed the same analysis for tarsal claw and mouthpart (stylet and labium) dimensions from a previous study (14) vs body length to test for agreement with power law scaling with body length, and isometric scaling(c = 1) in particular. Because scaling law fits to both mouthpart lengths vs body length did not include any adult data within the fit confidence intervals, we also performed fits to only the data for nymphs.

Previous research has found that the adhesive pad area, A_adh_, scales linearly with body mass for organisms over a wide range of taxa and body sizes (29). For comparison, isometric scaling predicts that A_adh_ ∝ L^2^ ∝ M^2/3^. We therefore tested whether either of these relationships hold for SLFs using published data for their arolium dimensions (14) to estimate the arolium area (S2 Appendix); we then performed scaling law fits to these data vs body mass using the methods described above. Because the arolium does not increase monotonically in size (i.e., it is smaller on average for adults than for 4th instars) we fitted only the values for nymphs.

## Results

### Instar determination

The results of our morphometric measurements are shown in Fig 2 along with clustering data using the GMM model for mass vs body length. (See full results in S3 Appendix.) The data were sorted into identical clusters using GMM clustering for mass vs length and using k-means on lengths only. The cluster centroids provide estimates for the mean body length and mass for each life stage, which we compare with earlier studies in Fig. 3. As can be seen in Fig. 3A and Table 1, the body lengths for each instar agreed closely for the two years studied here, but not with values from earlier studies. The body masses were lower for early instars for the 2022 data than for those in the 2021 data, but greater for 4^th^ instars.

**Fig 2.**
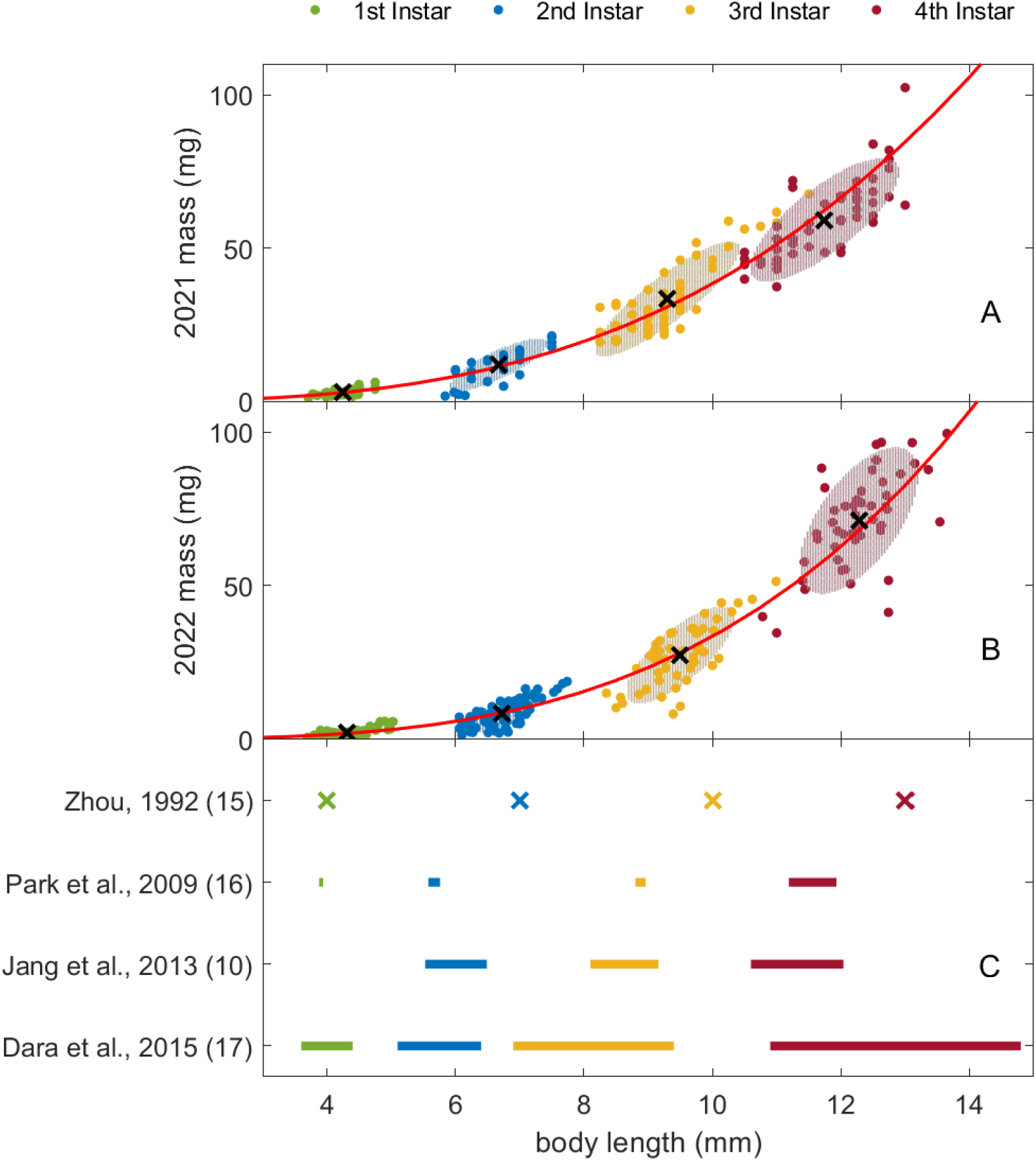
Clustering of spotted lanternfly nymph mass vs length data. Body mass vs length (filled circles) for (A) 2021 and (B) 2022 data; cluster centroids are shown as black x markers and red lines indicate scaling law fits to all data. Shaded ellipses show the 95% CI for each cluster based on the Mahalanobis distance; note that the shaded ellipses for some clusters are covered by datapoints. (C) Symbols and horizontal lines show the means and ranges of lengths, respectively, for each instar reported in previous studies (Table 1).

**Fig. 3.**
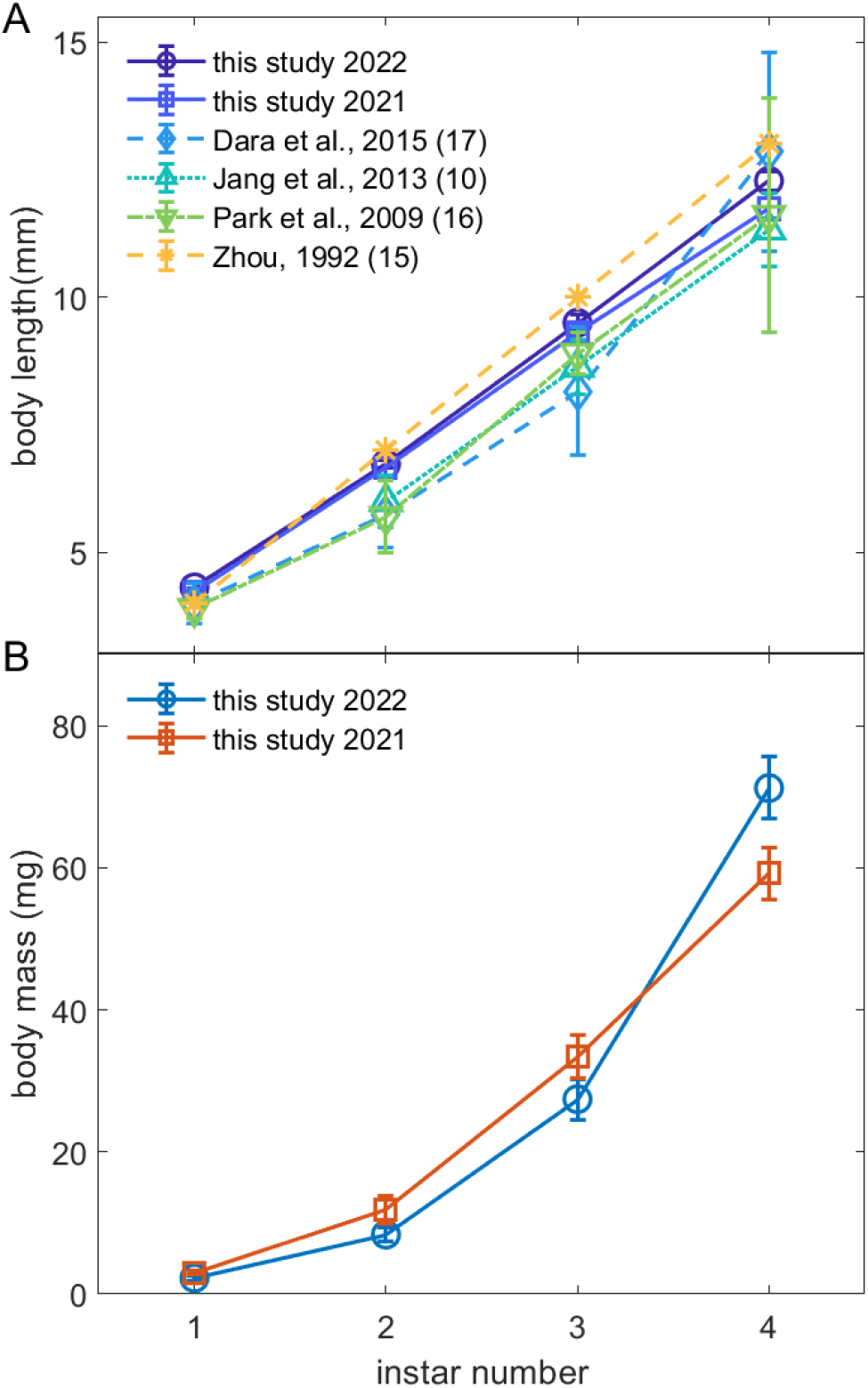
Comparison of body length and mass for each spotted lanternfly nymph stage from this study and previous research. (A) Spotted lanternfly nymph body length vs instar from this study and previous work (Table 1). (B) Spotted lanternfly nymph body mass vs instar from this study. Error bars are 95% CI for this study and the measures of variance given in Table 1. (Lines between datapoints show overall trends.)

Fig 4A show the results from testing Dyar’s Rule for SLF instar body length for this study and four previous studies. (See S4 Appendix for full results.) For every study except (10), body length was significantly related to instar number and the linear fit explained over 97% of total variance in the data in agreement with Dyar’s Rule. The fitted growth ratio for our 2022 data, G = 1.42 [1.25, 1.61], was consistent with that from 2021 and earlier studies (p > 0.95). (Fig 4B)

**Fig 4.**
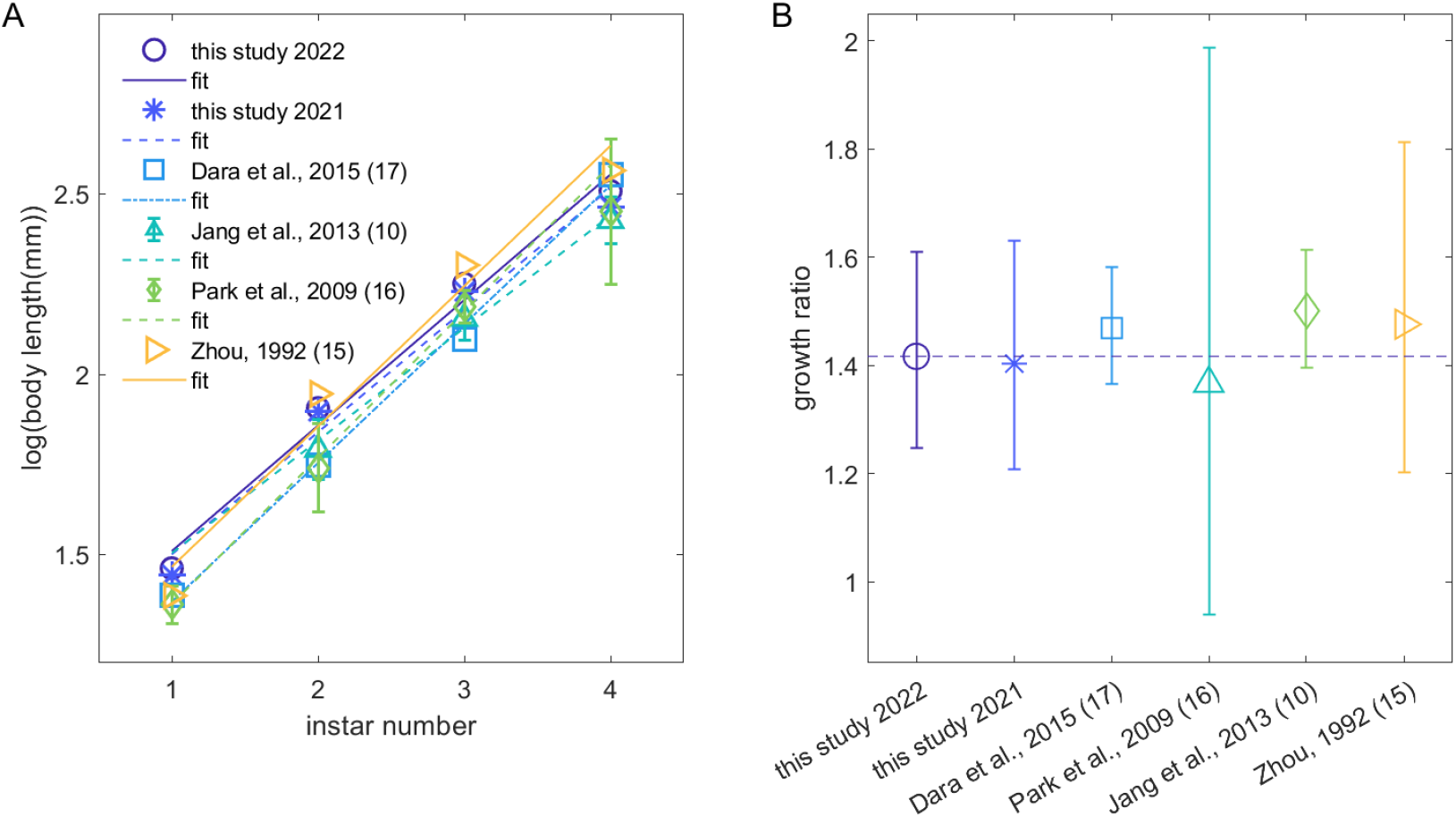
Results from Dyar’s Rule analysis of data from this study and previous research. (A) Plots of spotted lanternfly nymph log(body length) plotted vs estimated instar number as a test of Dyar’s rule. Markers show data from this study and three previous papers, while lines indicate linear fits. (B) Growth ratios, G, from the fits in (A); dashed line indicates the mean value for 2022 from this study. All error bars are 95% CI.

### Allometry

The SLF nymph body mass vs length data followed power law scaling with significantly different scaling exponents for the two years studied: 3.01 [2.94,3.09] for 2021 data and 3.45 [3.34, 3.56] for 2022 (Fig 2); the results for 2021 were thus consistent with isometric scaling while those for 2022 indicated positive allometry. (See S4 Appendix for full results for all scaling law fits.) Given the close agreement between instar length distributions in 2021 and 2022, we only used 2022 data in the analysis of foot and mouthpart data from (14). Fit results revealed that power law models accounted for a high percent of total variance for the footpart dimensions considered (77% and 62% for TCT and A_adh_, respectively). This also showed that SLF tarsal claw tip-to-tip distance (14) scales with body length over all SLF life stages with c = 1.33 [1.09, 1.56], indicating that this measure of the tarsal claws’ grasp displays positive allometry. (Fig 5A) SLF nymph arolium area depended on L with scaling exponent c_L_ = 2.49 [1.75, 3.23]; this corresponded to A_adh_ ∝ M ^0.89 ± 0.27^ (2021) and M ^0.73 ± 0.22^ (2022), in agreement with the dependence found for other taxa (Table S1, (29). (Fig 5B) By contrast, the fits shows that SLF nymph mouthpart dimensions (labium length, L_L_, and stylet length, L_S_) were only weakly correlated with body length (S4 Appendix, Fig 5B,C); i.e., power law scaling explains only 36% of the total variance in these data.

**Fig 5.**
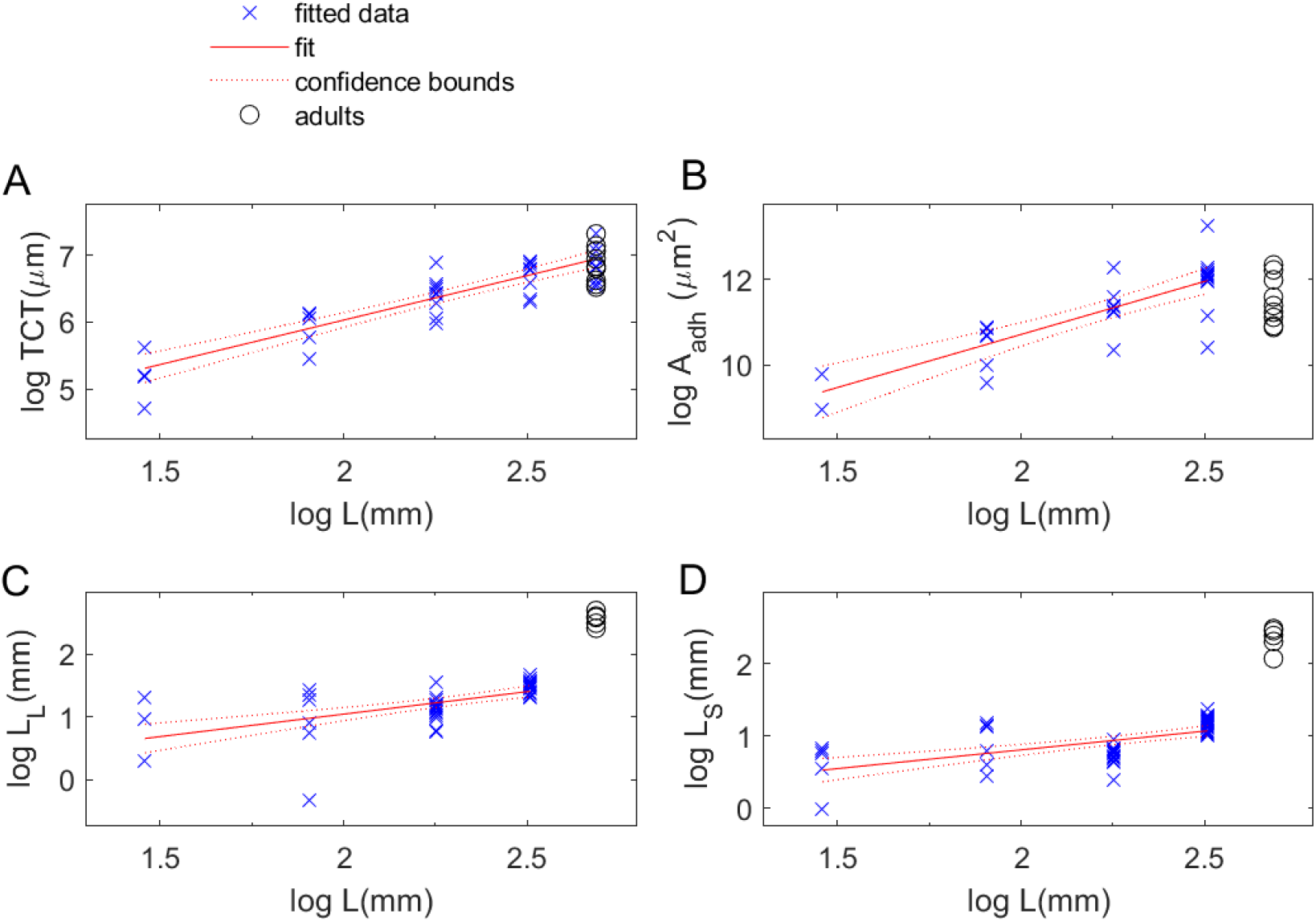
Plots and fits showing how spotted lanternfly foot and mouthpart data relate to body size measures. A) Plots of data and fit results for the dependence on log body length for (A) the log tarsal claw tip-to-tip width, TCT, (B) arolium area, A_adh_, (C) labium length, L_L_, and (D) stylet length, L_S_. Footpart and mouthpart morphometric data are from (14), while nymph body length and mass are from this study and adult mass and length from (27). (Fit lines and 95% confidence intervals from ordinary linear regression fits to power laws, as described in the main text.)

## Discussion

The results of this study lead to several conclusions. First, the distribution of measured data for body mass and length for 1^st^ to 3^rd^ instars were consistent with three non-overlapping clusters of data in the approximate size ranges expected for these life stages from previous studies. We consequently used two methods to assign these data to three clusters: 1) GMM on both mass and length data; 2) k-means clustering applied to length-only data. Both methods resulted in identical cluster assignments. This indicates that easy-to-perform specimen body length measurements and k-means clustering should be sufficient for instar determination. (In contrast to the distinct ranges found for 1^st^ to 3^rd^ instars, the measured body length and mass data ranges for 3rd and 4th instars overlapped in both years of our study. Due to the red coloration of 4th instar nymphs, however, this overlap in ranges does not cause a logistical challenge for properly categorizing these life stages.)

Practical applications of these findings include the indication that clustering methods with large datasets of spotted lanternfly body lengths should facilitate determining the likely developmental stage of individual specimens in future studies. The measured datasets and clustering code provided as supplemental materials enable other researchers to replicate and extend these results either to estimate the life stage of new specimens based on our data, or to perform clustering on their own measurements.

Second, we consider how measures of body size varied between the two years of our study and in comparison with previous research. Between the two years of our study, the length data were consistent for all nymphal life stages, but we found significant differences between 2021 and 2022 mean body masses at each life stage, ranging from -17% to +9%. These mass differences were not simply due to the timing of sampling: e.g., in 2022 only one 4^th^ instar had been recorded by the last week sampled in 2021. Additional data could help resolve whether environmental factors, time of first emergence, or other factors might influence the mass at different life stages.

The values for growth ratio between instars found using Dyar’s Rule and the approximate length ranges reported for 1st instars were consistent for all studies considered here. However, there was considerable variation amongst the body lengths found by different studies for 2^nd^-4^th^ instars. These differences could be due to a variety of factors, including variations in study design (e.g., different sampling methods used to collect different life stages and different reported summary statistics), different environmental conditions, or differences in population morphology. This diversity of reported values for instar sizes suggests that it would be interesting to compile and analyze morphometric data as a function of life stage for SLFs raised under a variety of circumstances to understand how much of this variation is due to sampling methods and how much to environmental conditions (location, diet, temperature etc.)

Third, these data also provide insights into the ontogenetic allometric scaling of spotted lanternfly nymph body metrics. Body mass scaled with body length for SLF nymphs with different exponents consistent with an approximately constant overall geometric shape (isometric scaling, 2021) or positive allometry (2022).

Fourth, these data can be used to create new syntheses of existing research for greater insight into the biology of these insects. For example, Kim et al. (30) hypothesized that earlier SLF instars should be more easily dislodged by wind than later nymphs due to their smaller arolia, an idea with implications for how dispersal and control should depend on life stage. However, SLF nymph arolium area (the morphometric measure relevant for adhesive strength) from (14) was found to scale with extreme positive allometry with body length and mas, in agreement with the scaling relationship found across taxa over 7 orders of magnitude of body weight (29). In combination with the finding of constant (i.e., size-independent) maximum adhesive stress between the arolium and surface for other insect adhesive pads (e.g., the pulvilli of *Coreus marginatus* (30) and the arolia of stick insects in (31)) across all life stages, this implies that SLF instars could have similar adhesive capabilities across life stages. The reported monotonic increase in SLF falling-climbing cycle period with advancing date of the year (7) could be due to factors other than arolium development, such as faster than isometric growth of tarsal claw grasp along with the detailed morphometric changes reported in (14): increased wrinkling of the arolia surface and a larger terminal sticky lip in adult SLFs relative to nymphs. These findings for adhesion are of especial interest because of the crucial role transportation plays in the dispersal of SLFs, which are known to travel long distances by clinging to vehicles and shipping containers (32).

By contrast, the analysis showed that the variation in the stylet and labium lengths was only weakly correlated with body length for SLF nymphs (14). This is consistent with the expectation that stylet length is correlated with preferred host plant tissue characteristics (33), as opposed to insect size, given reports from the literature indicate that SLF nymphs only feed on herbaceous and non-woody parts of plants (e.g., shoots, stems and leaves) while adults are able to feed on bark-covered trunks (7,9,16,17,34).

## Conclusion

We propose that body mass vs length curves can play a role similar to that of clinical growth charts, filling the current gap in metrics of SLF development. These measures can serve as a fitness benchmark for interpreting data from field studies and laboratory experiments to assess the impact of factors such as date of first emergence, molt schedule, temperature, diet, and geographic location. Furthermore, the successful fits to Dyar’s rule provide a measure of the growth ratio between successive instars, which might serve as an additional metric for comparing populations grown under different circumstances. Another potential use of these methods involves estimating the life stage of isolated SLF nymphs found in new locations as these insects expand their range. This information can play a valuable role in determining the stage of infestation, informing control efforts as well as providing data useful for tracking and modeling their spread. We therefore suggest that morphometrics of SLF nymphs be incorporated into ongoing studies when possible so as to provide a wide range of data for such applications going forward.

## Acknowledgements

We wish to thank Chengpei Li, Aaron Xu, Aidan Bannon, Simon Thill, and Sophie Frem for assistance in collecting specimens.

## S1 Appendix.

Sampling timeline for spotted lanternfly nymph body length and mass

**S1 Table.** Number of weeks over which each nymphal life stage was sampled.

|  | Number of weeks collected |  |
| --- | --- | --- |
| Life stage | 2021 | 2022 |
| 1st instar | 2 | 5 |
| 2nd instar | 4 | 5 |
| 3rd instar | 3 | 5 |
| 4th instar | 3 | 4 |

**S1 Fig.**
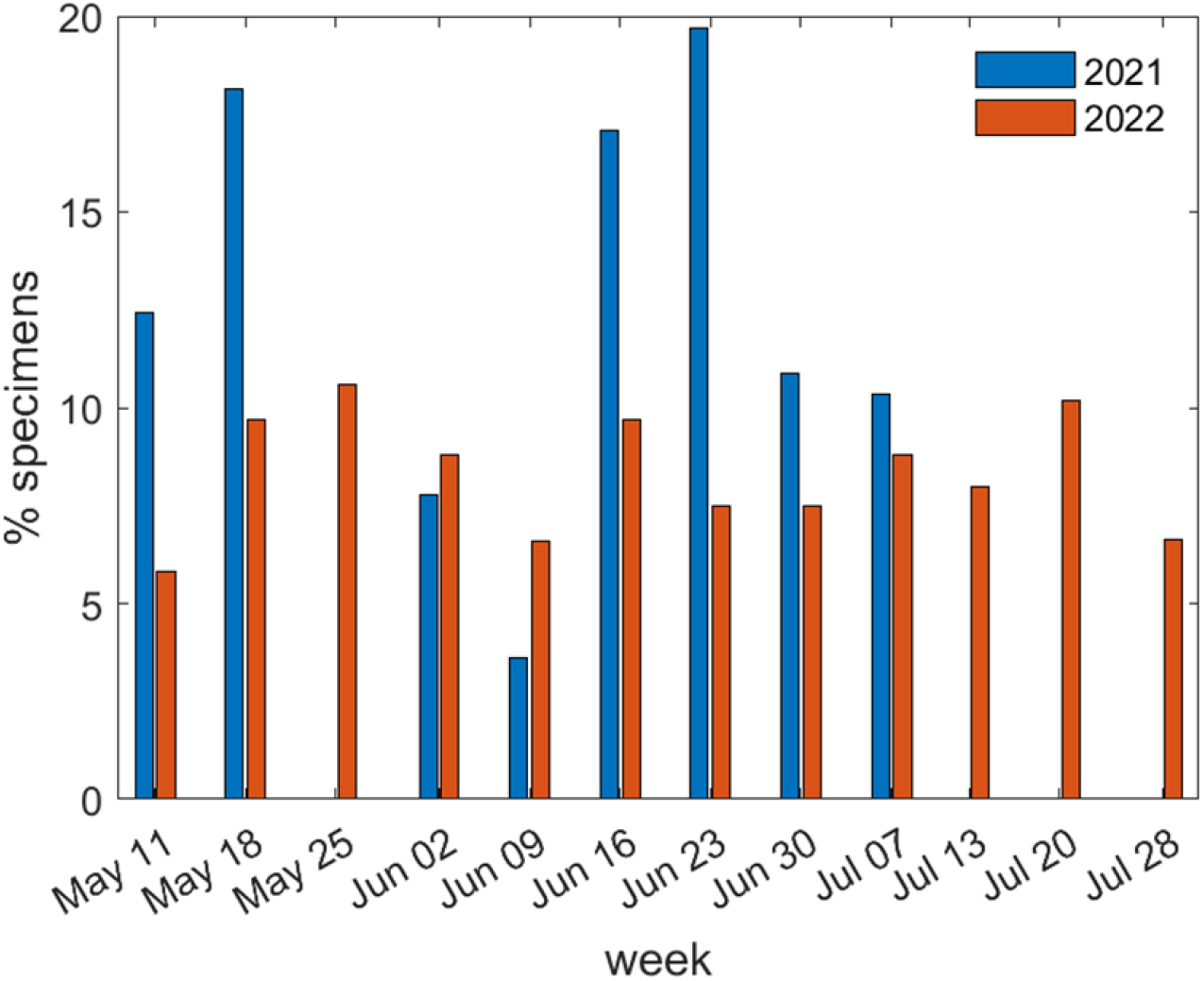
Sampling timeline for summer 2021 & 2022.

## S2 Appendix.

Spotted lanternfly arolium area

We used the data from Table 5 in (1) for spotted lanternfly arolium (foot adhesive pad) dimensions using the approximately triangular geometry defined in Fig 1 in ref. (1). This gives an estimated arolium area, *A*_*adh*_:

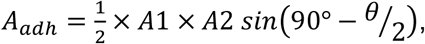

where the relevant arolium dimensions are defined in S2 Fig and S2 Table below.

**S2 Table.** Definitions of labels for arolium dimensions.

|  | Label in Fig 1 ref. (1) | Label in Table 5 ref. (1) |
| --- | --- | --- |
| maximum anterior width of the arolium | A1 | AAW |
| length of the lateral margin of the arolium | A2 | ASL |
| angle between the lateral margins of the arolium from the dorsal view | $\theta$ | AAG |

**S2 Fig.**
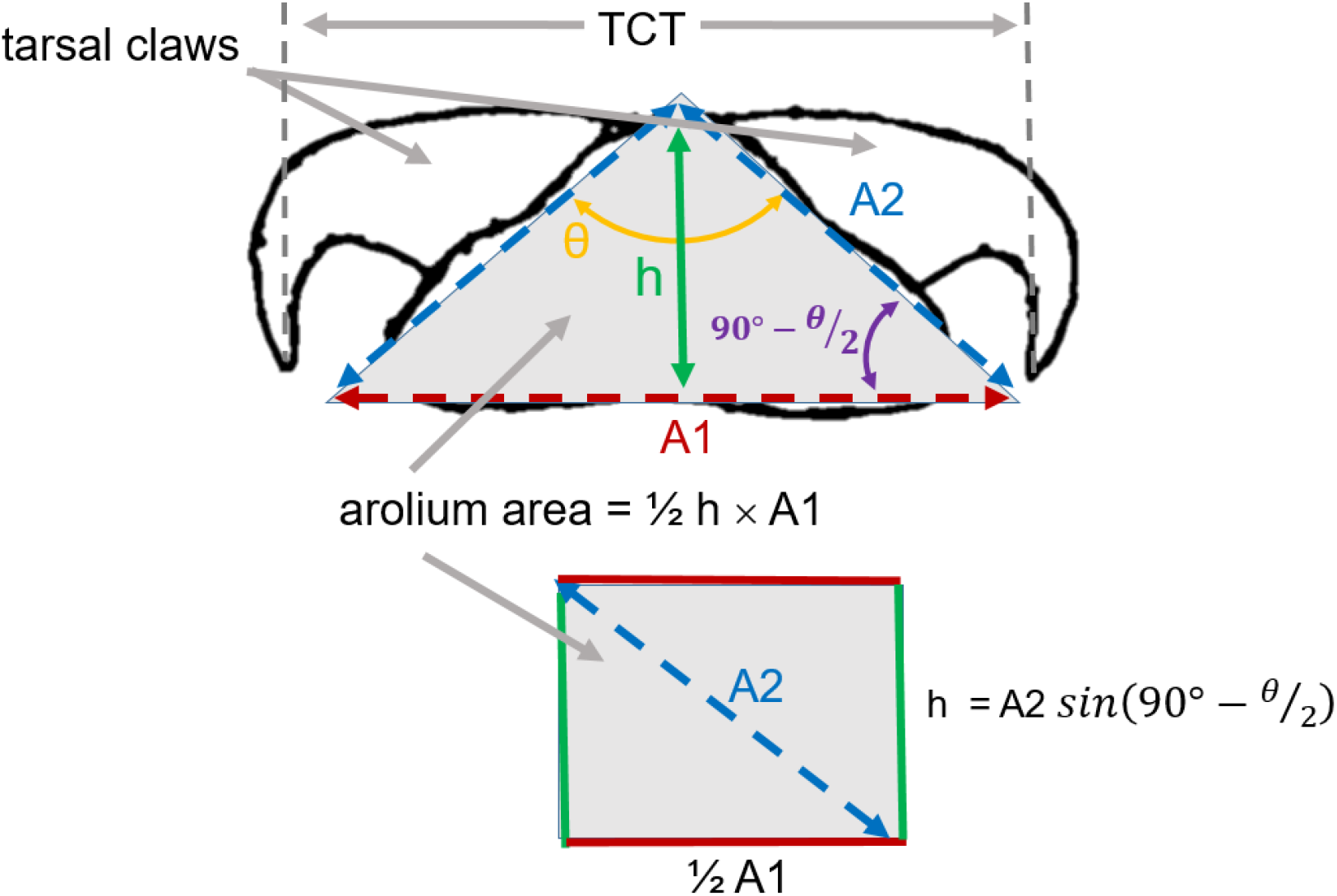
Approximately triangular geometry used to define the arolium’s dimensions. TCT = tarsal claw tip separation (adapted from Fig 1 in ref. (1)).

## S3 Appendix.

Clustering results

**Table S3.** Morphometric data for spotted lanternfly nymph mass and body length. The 4th instars were identified by coloration, while 1st, 2nd and 3rd instars were classified using fits of the mass vs length to a 3 component Gaussian mixture model (see Methods in main text for details). (N = number specimens. All values are given as mean ± SD.)

| Life stage | N<br>2022 | Body length<br>(mm)<br>2022 | Body mass<br>(mg)<br>2022 | N<br>2021 | Body length<br>(mm)<br>2021 | Body mass (mg)<br>2021 |
| --- | --- | --- | --- | --- | --- | --- |
| 1st instar | 49 | 4.32 ± 0.35 | 2.2 ± 1.4 | 54 | 4.24 ± 0.24 | 3.0 ± 1.2 |
| 2nd instar | 88 | 6.73 ± 0.38 | 8.4 ± 4.2 | 30 | 6.67 ± 0.48 | 12.0 ± 5.5 |
| 3rd instar | 49 | 9.50 ± 0.54 | 27.4 ± 10.2 | 61 | 9.30 ± 0.72 | 33.4 ± 12.0 |
| 4th instar | 40 | 12.28 ± .60 | 71.2 ± 15.4 | 49 | 11.74 ± .75 | 59.2 ± 13.0 |

## S4 Appendix.

Full fitting & data analysis results

**S4 Table.** Results from fits to Dyar’s Rule for data from this study and previously-published work. Fit to Dyar’s Rule: log L_j_ = j × log G + log L_0_, where j = instar number, L _j_ = mean body length of the jth instar, and G = growth ratio = L _j+1_ / L _j_ (dof = fit degrees of freedom. Note that when log L_0_ is not significantly different from 0, this simply means that L_0_= 1 and does not indicate a poor fit to the allometric scaling law.)

| Data | log L | t | p | G | t | p | R-squared | F | P | dof |
| --- | --- | --- | --- | --- | --- | --- | --- | --- | --- | --- |
| this study<br>2022 | $1.16 \pm 0.08$<br>[0.81, 1.51] | 14 | 0.005 | $1.42 \pm 0.04$<br>[1.25, 1.61] | 12 | 0.007 | 0.986 | 138 | 0.007 | 2 |
| this study<br>2021 | $1.16 \pm 0.10$<br>[0.75, 1.57] | 12 | 0.007 | $1.40 \pm 0.05$<br>[1.21, 1.63] | 9.7 | 0.01 | 0.979 | 95 | 0.01 | 2 |
| Dara et<br>al., 2015<br>(17) | $0.98 \pm 0.05$<br>[0.78, 1.18] | 21 | 0.002 | $1.47 \pm 0.02$<br>[1.37, 1.58] | 23 | 0.002 | 0.996 | 518 | 0.002 | 2 |
| Jang et al.,<br>2013 (10) | $1.19 \pm 0.10$<br>[-0.02, 2.41] | 12 | 0.05 | $1.37 \pm 0.04$<br>[0.94, 1.99] | 11 | 0.06 | 0.991 | 112 | 0.06 | 1 |
| Park et al.,<br>2009 (16) | $0.96 \pm 0.04$<br>[0.78, 1.13] | 24 | 0.002 | $1.50 \pm 0.03$<br>[1.40, 1.61] | 24 | 0.002 | 0.997 | 583 | 0.002 | 2 |
| Zhou et<br>al., 1992<br>(15) | $1.08 \pm 0.13$<br>[0.52, 1.64] | 8.3 | 0.014 | $1.478 \pm 0.07$<br>[1.20, 1.81] | 8.2 | 0.01 | 0.971 | 67 | 0.01 | 2 |

**S5 Table.** Full results from allometric fits to y = a + c log x. Log-transformed values for the spotted lantemfly body mass = M (mg), tarsal claw tip distance = TCT (μm), nymph labium length = L_L_ (mm) and stylet length = L_s_(mm) from (14) were fitted vs log-transformed body length = L (mm) from this study and (27). Log arolium area = A_adh_, (micrometer^2^) computed from data in (14) was fitted to log M from this study and (27). Fit parameters are intercept = a, and slope, c = scaling exponent. For M vs L fits, because some specimens has residuals that were outliers due to low mass (as determined using MATLAB *isoutlier*, < 3 median absolute deviation threshold), with disproportionately fiat abdomens presumably due to hunger), we conducted fits with and without excluding outliers (5 for 2021 and 13 for 2022 data); scaling exponents for these fits differed by 1% so we report only results excluding outliers here.

| Data | Intercept, $a$ | $t$ | $p$ | scaling exponent, $c$ | $t$ | $p$ | R-squared | F | P | dof |
| --- | --- | --- | --- | --- | --- | --- | --- | --- | --- | --- |
| $M$ vs $L$<br>(2022) | $-4.42 \pm 0.11$<br>[-4.65, -4.20] | -39 | < 0.001 | $3.45 \pm 0.07$<br>[3.34, 3.56] | 48 | < 0.001 | 0.948 | 2323 | < 0.001 | 211 |
| $M$ vs $L$<br>(2021) | $-3.29 \pm 0.08$<br>[-3.44, -3.13] | -43 | < 0.001 | $3.01 \pm 0.04$<br>[2.94, 3.09] | 80 | < 0.001 | 0.972 | 6467 | < 0.001 | 187 |
| $TCT$ vs. $L$ | $3.38 \pm 0.27$<br>[2.83, 3.92] | 13 | < 0.001 | $1.33 \pm 0.12$<br>[1.09, 1.56] | 11.4 | < 0.001 | 0.766 | 131 | < 0.001 | 40 |
| $A_{aoh}$ vs $L$ | $5.74 \pm 0.81$<br>[4.08, 7.40] | 7.1 | < 0.001 | $2.49 \pm 0.36$<br>[1.75, 3.23] | 6.9 | < 0.001 | 0.620 | 47 | < 0.001 | 29 |
| $L_S$ vs $L$ | $-0.23 \pm 0.21$<br>[-0.65, 0.20] | -1.1 | 0.29 | $0.52 \pm 0.09$<br>[0.33, 0.70] | 5.6 | < 0.001 | 0.357 | 31 | < 0.001 | 56 |
| $L_L$ vs $L$ | $-0.39 \pm 0.30$<br>[-0.99, 0.21] | -1.3 | 0.19 | $0.72 \pm 0.13$<br>[0.46, 0.98] | 5.5 | < 0.001 | 0.363 | 31 | < 0.001 | 54 |

## References

1. Murman K, Setliff GP, Pugh CV, Toolan MJ, Canlas I, Cannon S, et al. Distribution, Survival, and Development of Spotted Lanternfly on Host Plants Found in North America. Environ Entomol. 2020 Dec 14;49(6):1270–81.

2. Liu H. Oviposition Substrate Selection, Egg Mass Characteristics, Host Preference, and Life History of the Spotted Lanternfly (Hemiptera: Fulgoridae) in North America. Environmental Entomology. 2019 Dec 2;48(6):1452–68.

3. Calvin DD, Keller J, Rost J, Walsh B, Biddinger D, Hoover K, et al. Spotted Lanternfly (Hemiptera: Fulgoridae) Nymphal Dispersion Patterns and Their Influence on Field Experiments. Environmental Entomology. 2021 Dec 1;50(6):1490–504.

4. Keller JA, Johnson AE, Uyi O, Wurzbacher S, Long D, Hoover K. Dispersal of Lycorma delicatula (Hemiptera: Fulgoridae) Nymphs Through Contiguous, Deciduous Forest. Environmental Entomology. 2020 Oct 17;49(5):1012–8.

5. Cooperband M, Murman K, Cannon S, Abreu L, Wallace M, Trepanowski N, et al. Dispersal and host preference of marked and released spotted lanternfly. Otis laboratory 2018 annual report. United States Department of Agriculture Buzzards Bay, MA; 2019. 60–61 p.

6. Cooperband MF, Wickham J, Cleary K, Spichiger SE, Zhang L, Baker J, et al. Discovery of Three Kairomones in Relation to Trap and Lure Development for Spotted Lanternfly (Hemiptera: Fulgoridae). J Econ Entomol. 2019 Mar 21;112(2):671–82.

7. Kim JG, Lee EH, Seo YM, Kim NY. Cyclic Behavior of Lycorma delicatula (Insecta: Hemiptera: Fulgoridae) on Host Plants. J Insect Behav. 2011 Jun 8;24(6):423.

8. Nixon LJ, Ludwick DC, Leskey TC. Horizontal and vertical dispersal capacity and effects of fluorescent marking on Lycorma delicatula nymphs and adults. Entomologia Experimentalis et Applicata. 2021;169(2):219–26.

9. Leach H, Leach A. Seasonal phenology and activity of spotted lanternfly (Lycorma delicatula) in eastern US vineyards. Journal of Pest Science. 2020;93:1215–24.

10. Jang Y, An HG, Kim H, Kim KH. Spectral preferences of Lycorma delicatula (Hemiptera: Fulgoridae). Entomological Research. 2013;43(2):115–22.

11. Moon SR, Cho SR, Jeong JW, Shin YH, Yang JO, Ahn KS, et al. Attraction response of spot clothing wax cicada, Lycorma delicatula (Hemiptera: Fulgoridae) to spearmint oil. Journal of the Korean Society for Applied Biological Chemistry. 2011;54(4):558–67.

12. Desko M, Schiebel C, Silverman S, Bickel J, Felton K, Chandler JL. The Probability of Spotted Lanternfly, Lycorma delicatula (Hemiptera: Fulgoridae), Escape Differs Among Life Stages and Between Two Trapping Techniques Commonly Used By Landowners, Sticky Bands and Duct Tape. The Great Lakes Entomologist. 2020;53(2):10.

13. Wang RR, Liu JJ, Li XY, Liang AP, Bourgoin T. Relating antennal sensilla diversity and possible species behaviour in the planthopper pest Lycorma delicatula (Hemiptera: Fulgoromorpha: Fulgoridae). PLoS One [Internet]. 2018 Mar 27 [cited 2020 Jun 16];13(3). Available from: https://www.ncbi.nlm.nih.gov/pmc/articles/PMC5871016/

14. Avanesyan A, Maugel TK, Lamp WO. External morphology and developmental changes of tarsal tips and mouthparts of the invasive spotted lanternfly, Lycorma delicatula (Hemiptera: Fulgoridae). PLoS One [Internet]. 2019 Dec 26 [cited 2020 Jun 16];14(12). Available from: https://www.ncbi.nlm.nih.gov/pmc/articles/PMC6932783/

15. Zhou JX. Lycorma delicatula (White)(Homoptera: Fulgoridae). Forest insects of China, 2nd ed Chinese Forestry Publishing House, Beijing, China. 1992;169–70.

16. Park JD, Kim MY, Lee SG, Shin SC, Kim JH, Park IK. Biological Characteristics of Lycorma delicatula and the Control Effects of Some Insecticides. Korean journal of applied entomology. 2009;48(1):53–7.

17. Dara SK, Barringer L, Arthurs SP. Lycorma delicatula (Hemiptera: Fulgoridae): A New Invasive Pest in the United States. J Integr Pest Manag. 2015 Mar 1;6(1).

18. Daly HV. Insect morphometrics. Annual review of entomology. 1985;30(1):415–38.

19. Uyi O, Keller JA, Johnson A, Long D, Walsh B, Hoover K. Spotted Lanternfly (Hemiptera: Fulgoridae) Can Complete Development and Reproduce Without Access to the Preferred Host, Ailanthus altissima. Environmental Entomology. 2020 Oct 17;49(5):1185–90.

20. Uyi O, Keller JA, Swackhamer E, Hoover K. Performance and host association of spotted lanternfly (Lycorma delicatula) among common woody ornamentals. Sci Rep. 2021 Aug 4;11(1):15774.

21. Kreitman D, Keena MA, Nielsen AL, Hamilton G. Effects of Temperature on Development and Survival of Nymphal Lycorma delicatula (Hemiptera: Fulgoridae). Environmental Entomology. 2021 Feb 1;50(1):183–91.

22. Chen Y, Seybold SJ. Application of a Frequency Distribution Method for Determining Instars of the Beet Armyworm (Lepidoptera: Noctuidae) From Widths of Cast Head Capsules. Journal of Economic Entomology. 2013 Apr 1;106(2):800–6.

23. Rinehart LF, Rasnitsyn AP, Lucas SG, Heckert AB. Instar sizes and growth in the Middle Permian monuran Dasyleptus brongniarti (Insecta: Machilida: Dasyleptidae). New Mex Mus Nat Hist Sci Bull. 2005;30:270–2.

24. Sohlström EH, Marian L, Barnes AD, Haneda NF, Scheu S, Rall BC, et al. Applying generalized allometric regressions to predict live body mass of tropical and temperate arthropods. Ecology and evolution. 2018;8(24):12737–49.

25. Pennsylvania Department of Agriculture. Spotted Lanternfly [Internet]. Department of Agriculture. [cited 2022 Feb 10]. Available from: https://www.agriculture.pa.gov/Plants_Land_Water/PlantIndustry/Entomology/spotted_lanternfly/Pages/default.aspx

26. Schneider CA, Rasband WS, Eliceiri KW. NIH Image to ImageJ: 25 years of image analysis. Nature Methods. 2012 Jul;9(7):671–5.

27. Frantsevich L, Ji A, Dai Z, Wang J, Frantsevich L, Gorb SN. Adhesive properties of the arolium of a lantern-fly, Lycorma delicatula (Auchenorrhyncha, Fulgoridae). Journal of Insect Physiology. 2008 May 1;54(5):818–27.

28. Dyar HG. The number of molts of lepidopterous larvae. Psyche. 1890;5(175–176):420–2.

29. Labonte D, Clemente CJ, Dittrich A, Kuo CY, Crosby AJ, Irschick DJ, et al. Extreme positive allometry of animal adhesive pads and the size limits of adhesion-based climbing. PNAS. 2016 Feb 2;113(5):1297–302.

30. Gorb SN, Gorb EV. Ontogenesis of the attachment ability in the bug Coreus marginatus (Heteroptera, Insecta). Journal of Experimental Biology. 2004 Aug 1;207(17):2917–24.

31. Labonte D, Struecker MY, Birn-Jeffery AV, Federle W. Shear-sensitive adhesion enables size-independent adhesive performance in stick insects. Proceedings of the Royal Society B: Biological Sciences. 2019 Oct 23;286(1913):20191327.

32. Urban JM. Perspective: shedding light on spotted lanternfly impacts in the USA. Pest Management Science. 2020;76(1):10–7.

33. Leopold RA, Freeman TP, Buckner JS, Nelson DR. Mouthpart morphology and stylet penetration of host plants by the glassy-winged sharpshooter, Homalodisca coagulata, (Homoptera: Cicadellidae). Arthropod Structure & Development. 2003 Oct 1;32(2):189–99.

34. Barringer L, Ciafré CM. Worldwide Feeding Host Plants of Spotted Lanternfly, With Significant Additions From North America. Environmental Entomology. 2020 Oct 17;49(5):999–1011.

## References

1. Avanesyan A, Maugel TK, Lamp WO. External morphology and developmental changes of tarsal tips and mouthparts of the invasive spotted lanternfly, Lycorma delicatula (Hemiptera: Fulgoridae). PLoS One [Internet]. 2019 Dec 26 [cited 2020 Jun 16];14(12). Available from: https://www.ncbi.nlm.nih.gov/pmc/articles/PMC6932783/

2. Frantsevich L, Ji A, Dai Z, Wang J, Frantsevich L, Gorb SN. Adhesive properties of the arolium of a lantern-fly, Lycorma delicatula (Auchenorrhyncha, Fulgoridae). Journal of Insect Physiology. 2008 May 1;54(5):818–27.

